# Mycophenolate mofetil reduces the branching of microglial processes

**DOI:** 10.1101/2025.11.12.687812

**Authors:** Rin-ichiro Teruya, Kentaro Ueda, Takumi Taketomi, Takushi Yamamoto, Naoki Yamashita, Hana Konno, Fuminori Tsuruta

## Abstract

Microglia, the resident immune cells in the central nervous system, play important roles not only in immune response but also in neurogenesis, synaptogenesis, and neural circuit formation. Microglia also surveil the brain environment via elongation and retraction of their processes. Previously, we found that the purine salvage pathway is involved in the regulation of morphology and dynamics of the microglial cell line BV2. Here, we show that intraperitoneal administration of mycophenolate mofetil (MMF), an inosine monophosphate dehydrogenase (IMPDH) inhibitor, reduces microglial branching during postnatal development. Imaging mass spectrometry analysis revealed that MMF administration decreases guanosine nucleotides in the brain. Interestingly, despite the essential role of guanosine nucleotides in cellular proliferation, MMF administration did not significantly affect microglial proliferation. On the other hand, MMF administration attenuated the level of GTP-bound forms of RhoA and Rac1 small GTPases. Notably, MMF administration decreased the number of branches, while process length remained unaffected. Since microglial branching affects microglial complexity and diversity, our findings suggest that guanosine nucleotide production is essential for generating proper microglial diversity.

## Introduction

Microglia are primary immune cells in the brain and play a key role in regulating inflammatory responses. Recent studies have shown that microglia originate from erythro-myeloid progenitors generated in the yolk sac during early embryonic stages and migrate into the brain primordium during mid-embryonic stages (1). After migration, these cells differentiate into mature microglia, which contribute to brain homeostasis through phagocytosis of dead cells, removal of foreign substances, regulation of synaptogenesis, and maintenance of central nervous system (CNS) border regions (2). Notably, microglia show a wide variety of gene expression patterns during the early postnatal stages (3–6). In these stages, microglia exhibit morphological transformation, changing from a relatively simple morphology with few processes to a complex, ramified morphology characterized by numerous branches. However, the detailed mechanisms underlying the regulation of microglial morphology remain poorly understood.

Previously, we have reported a link between the morphological changes of microglia and purine metabolism. We initially found that stimulation with hypoxanthine promotes process elongation in the microglial cell line BV2 (7). Hypoxanthine is an intermediate product in purine metabolism and serves as the substrate for the salvage pathway. In this pathway, hypoxanthine is catalyzed by hypoxanthine-guanine phosphoribosyltransferase (HPRT) and is converted to inosine monophosphate (IMP). Whereas, in the degradation pathway, xanthine oxidase (XO) converts hypoxanthine to xanthine, which is then further converted to uric acid. Previously, we have reported that intraperitoneal administration of allopurinol, an inhibitor of XO, promotes process extension and branching in microglia in vivo (8). Because treatment with allopurinol leads to the accumulation of hypoxanthine in cells (7), hypoxanthine appears to be a key mediator regulating microglial morphology. On the other hand, IMP, which is the product of the other pathway utilizing hypoxanthine as a substrate, is a critical purine metabolic intermediate produced through both the de novo and salvage pathways, serving as a precursor for guanosine nucleotides. IMP is converted to xanthine monophosphate (XMP) by inosine monophosphate dehydrogenase (IMPDH). XMP is subsequently metabolized to guanosine monophosphate (GMP). Thus, IMPDH is considered a key metabolic enzyme in the biosynthesis of guanosine nucleotides, including guanosine diphosphate (GDP) and guanosine triphosphate (GTP).

Here, we report that pharmacological inhibition of IMPDH reduces the levels of guanosine nucleotides in the brain and the amount of GTP-bound forms of small GTPase proteins. We also found that IMPDH suppression reduces the complexity of microglial morphology in vivo. These data suggest that IMPDH plays a crucial role in regulating morphological changes in microglia. Thus, our findings may shed light on a novel perspective regarding microglial heterogeneity and diversity during early postnatal development.

## Results

### MMF administration decreases the guanosine nucleotides levels in the brain

Because IMP is a crucial metabolite underlying production of guanosine nucleotide in both de novo and salvage pathways (Fig 1A), we focus on the necessity of IMPDH for the production of guanosine nucleotides in vivo. To do this, we intraperitoneally administered the IMPDH inhibitor, mycophenolate mofetil (MMF), to CX3CR1^GFP/+^ mice daily from postnatal day 6 (P6) to P14, and the relative abundance of guanosine nucleotides in the brain was then quantified using iMScope QT imaging mass spectrometry (IMS). IMS analysis detected the GMP ion at m/z 362.049, guanosine diphosphate (GDP) ion at m/z 442.012 and guanosine triphosphate (GTP) ion at m/z 521,98 (Fig. 1B and 1C). Due to the inherent instability of GTP, its signal was not suitable for precise quantification, whereas GMP and GDP were detected. We therefore used the m/z signals for reconstructing MS images of guanosine nucleotides using IMAGEREVEAL MS software (Fig. 1D). Quantitative analysis of these images revealed that MMF administration significantly decreased the guanosine nucleotide levels, particularly GMP and GDP, in the brain (Fig 1E to 1G).

**Figure 1.**
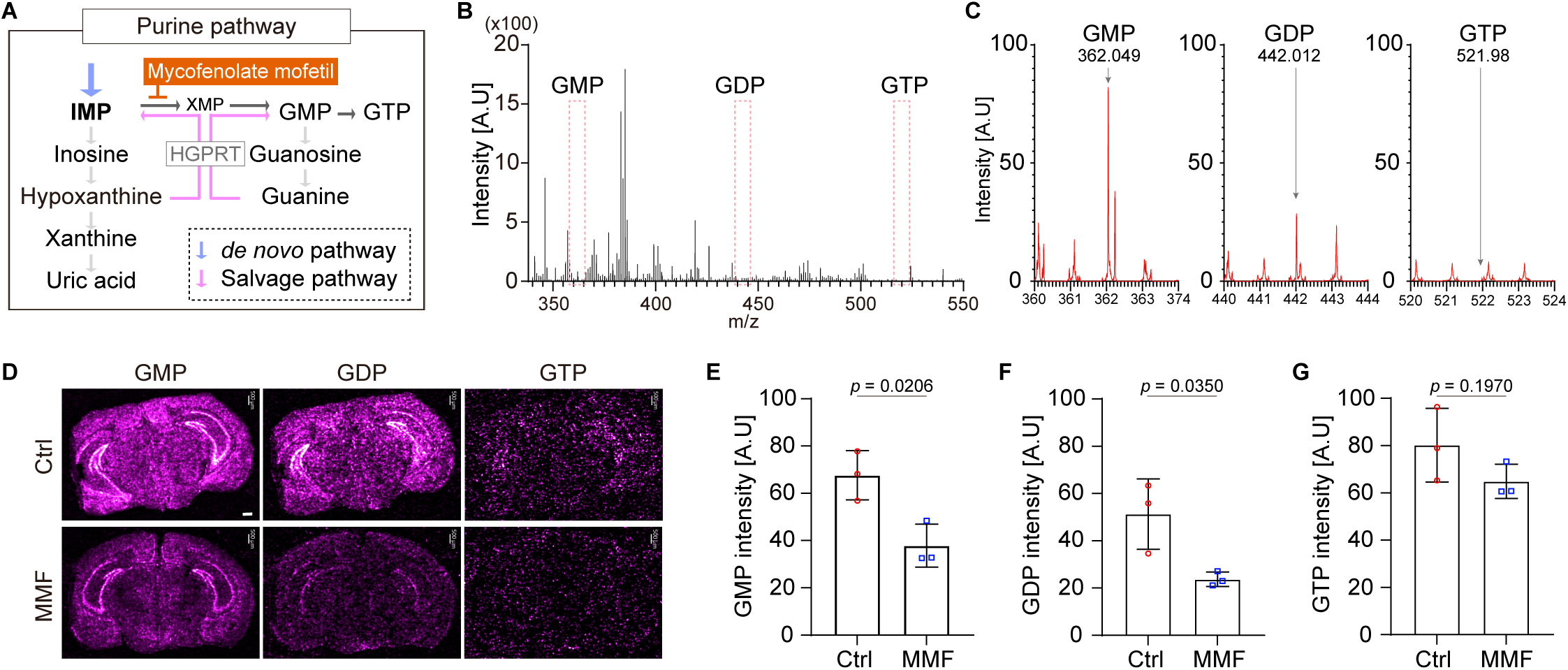
MMF administration decreases the guanosine nucleotides levels in the brain. **(A)** The de novo and salvage purine synthesis pathways. **(B)** Representative mass spectrum of mouse brain section (P14, WT) obtained from IMScope QT in negative ion mode. The signals were acquired at a m/z range of 340–550. **(C)** The expand insets of GMP (m/z = 360-374), GDP (m/z = 440-444) and GTP (m/z = 520-524) from the mass spectrum in B. **(D)** MMF (60 mg/kg) was administered intraperitoneally every 24 h from P6 to P14. IMS images obtained from P14 mice brain. The ion images show localization of GMP (m/z = 362.049), GDP (m/z = 442.012) or GTP (m/z = 521.98). Scale bar = 500 µm **(E to G)** The quantification of guanosine nucleotide signal intensity lelvels. The graphs show the average intensity of all dots in each section. One brain section was obtained from each individual each mouse. Ctrl; n = 3 sections, MMF; n = 3 sections, mean ± SEM, student’s *t*-test.

### MMF has little effect on microglial proliferation

MMF is used as an immunosuppressive agent that inhibits lymphocyte proliferation (9). Because microglia exhibit rapidly clonal expansion during early developmental stages (10), we investigated whether MMF affects microglial proliferation, similarly to lymphocytes. To do this, we analyzed the microglial density in vivo by administrating MMF to *Cx3cr1^GFP/+^* mice from P6 daily and collecting brains at P9 (the midpoint of microglial expansion) and P14 (the general terminal point of microglial expansion) (10,11) (Fig 2A). In contrast to the well-known effects of MMF on lymphocytes, our results show that the number of MMF-treated microglia is consistent with microglia without MMF administration (Fig 2B to 2E). These results suggest that MMF administration has little effect on microglial proliferation during the postnatal stages at least in this system.

**Figure 2.**
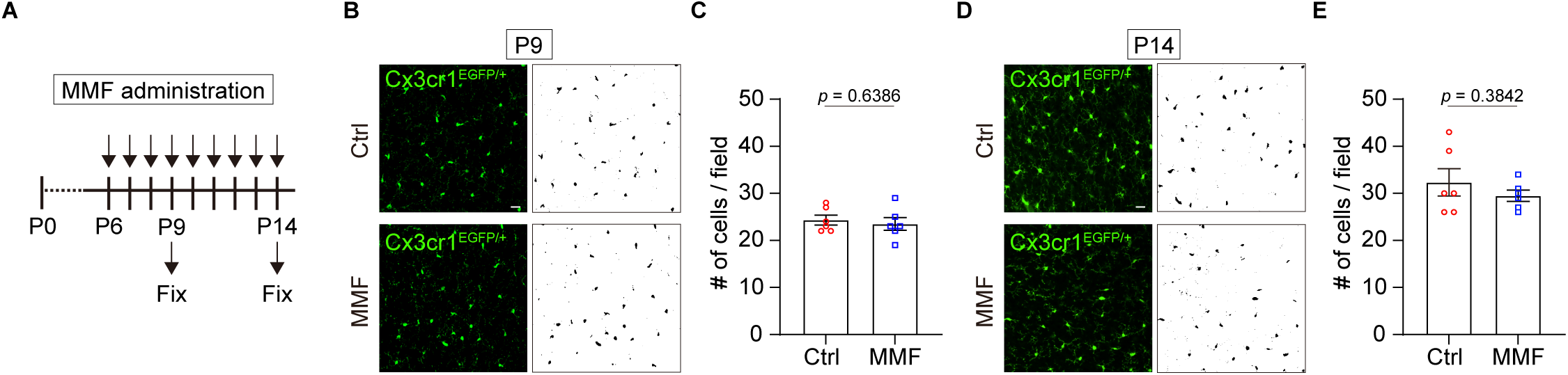
MMF has little effect on microglial proliferation. **(A)** Schematic of the experimental timeline. *Cx3cr1^GFP/+^* mice were administrated MMF (60 mg/kg) or a vehicle control daily from P6. Brains were collected for analysis at P9 and P14. **(B)** Immunostainings showing the expression of GFP in *Cx3cr1^GFP/+^* male mice at P9. The right panels show the binary images converted from the left panels. Scale bar = 20 µm. **(C)** Quantification of the number of microglia in B. One brain section was obtained from each individual mouse. Field = 1.03×10^5^ µm^2^. Ctrl; n = 3 sections, MMF; n = 3 sections, mean ± SEM, student’s *t*-test. **(D)** Immunostainings showing the expression of GFP in *Cx3cr1^GFP/+^* male mice in P14. The right panels show the binary images converted from the left panels. Scale bar = 20 µm. **(E)** Quantification of the number of microglia in D. One brain section was obtained from each individual mouse. Field = 1.03×10^5^ µm^2^. Ctrl; n = 3 sections, MMF; n = 3 sections, mean ± SEM, student’s *t*-test.

### Treatment with MMF suppresses morphological changes of BV2 cells

Guanosine nucleotides are essential for the regulation of small GTPase proteins, such as Rac1 and RhoA. These proteins are activated upon binding to GTP and subsequently regulate actin cytoskeletal dynamics, leading to changes in cell shape (Fig 3A). Since MMF treatment affects the pool of guanosine nucleotides (see Fig.1), we hypothesized that the activation of these small GTPases is reduced after MMF stimulation. To examine this, we performed the glutathione S-transferase (GST) -pull down assay in BV2 cells using recombinant GST-Pak CRIB domain (GST-CRIB) and GST-Rhotekin RBD domain (GST-RBD) (12,13). This assay revealed that the GTP-bound forms of Rac1 and RhoA were reduced after MMF treatment (Fig. 3B). Since the dynamic extension and retraction of microglial processes are regulated by these small GTPase proteins, we next investigated whether the MMF-induced reduction in GTP-bound forms would attenuate morphological dynamics of BV2 cells in an in vitro culture system (Fig. 3C). Time-lapse analysis revealed that control BV2 cells were highly dynamic, constantly transitioning between rounded and polarized morphologies. In contrast, MMF treatment suppressed these morphological changes, with cell shape maintaining a relatively stable throughout the imaging period (Fig. 3D). Notably, the variation in measurement values indicates that MMF treatment significantly reduced fluctuations in cell length and area (Fig 3E to 3H). These data suggest that MMF treatment impairs morphological dynamics of the BV2 microglial cell line.

**Figure 3.**
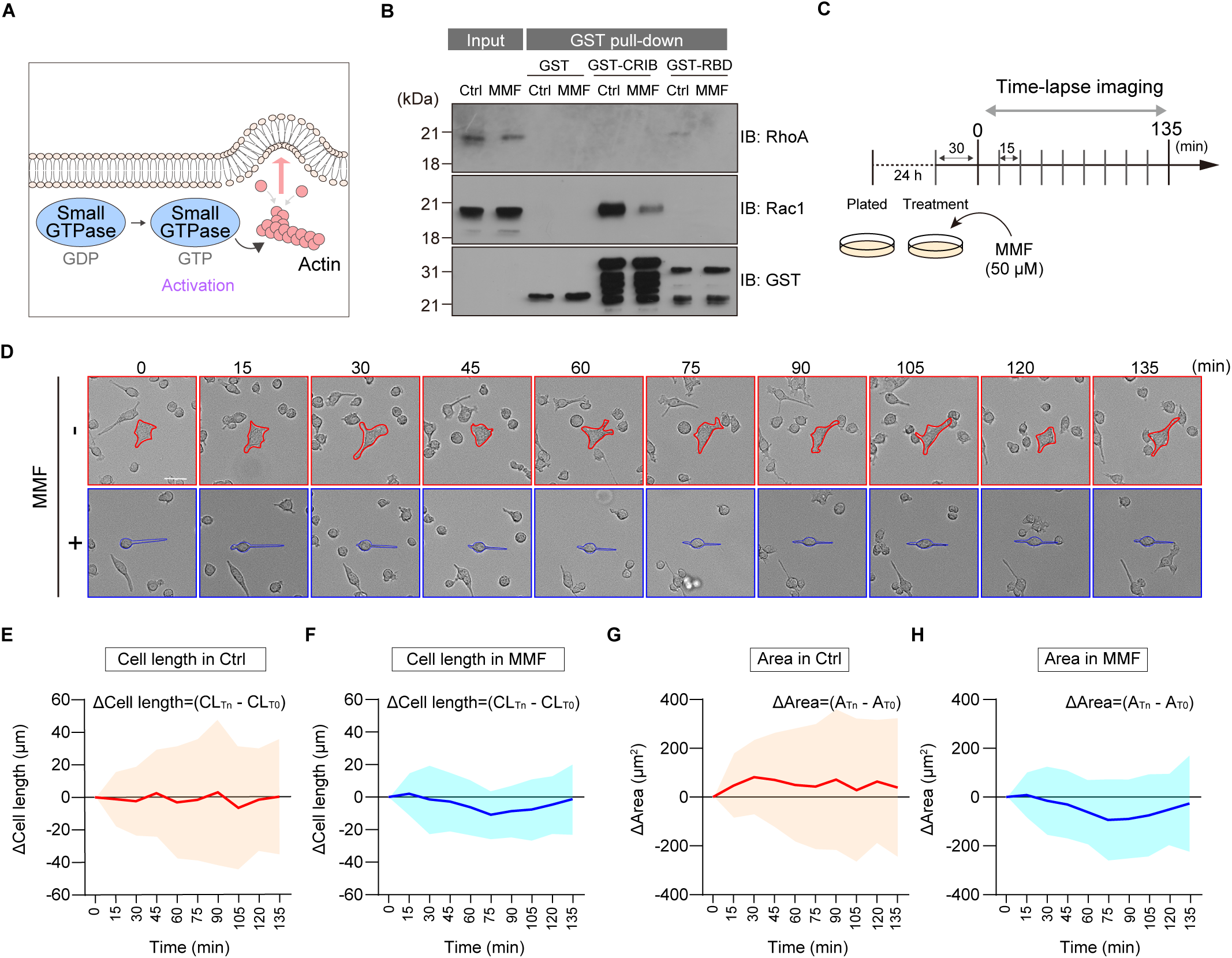
Treatment with MMF suppresses morphological changes of BV2 cells. **(A)** Schematic model of the small GTPase involved in cell morphology. **(B)** GST pull-down assay using BV2 cell lysates with or without MMF treatment (1 µM). Precipitated samples were subjected to immunoblot analysis and to detect GTP-bound forms of RhoA and Rac1. **(C)** Schematic of the live-cell imaging protocol. BV2 cells were pre-treated with vehicle (Ctrl) or MMF (1 µM) for 30 minutes, after which time-lapse images were acquired every 15 minutes for a total of 135 minutes. **(D)** Representative time-lapse images of BV2 cells treated with vehicle or MMF at the indicated time points. Scale bar = 20 µm. **(E-H)** Quantification of morphological dynamics over time using Fiji ImageJ. (E,F) graphs showing the change in Δ Cell length (CL_Tn_-CL_T0_) for individual Ctrl (E) or MMF treatment (F), (G,H) graphs showing the change in Δ Area (A_Tn_-A_T0_) for individual Ctrl (G) or MMF treatment (H), Ctrl ; n = 30 cells, MMF ; n = 30 cells. The cell length is defined by selecting the two points with the maximum distance.

### MMF treatment reduces the clustering of Rac1 in microglia

To confirm that our in vitro findings are applicable to an in vivo context, we next assessed the activation of small GTPase proteins using whole-brain lysates from MMF-injected mice. Consistent with our experiments in BV2 cells, a GST-pull down assay showed a significant reduction in the GTP-bound forms of both Rac1 and RhoA after MMF administration (Fig. 4A). Since we cannot rule out the possibility that intraperitoneal administration of MMF affects not only microglia but also other cells, such as neurons and astrocytes, we directly observed the subcellular localization of these proteins specifically within microglia using immunohistochemistry in *Cx3cr1^GFP/+^* mice. Because we were unable to detect endogenous RhoA signals due to an issue with antibody reactivity, we focused on Rac1 staining in this experiment. While we did not observe a diffuse cytoplasmic signal for Rac1, high-resolution imaging suggested that Rac1 is clustered in microglia (Fig. 4B and 4C). Because these signals align with the previous observation that Cdc42 and F-actin exhibit clustering structures involved in cytoskeletal rearrangement in microglia (14), we thought that these Rac1 clusters play a similar role in regulating microglial process dynamics. Then, we quantified the number of microglia with or without these clusters. Consequently, the number of microglia with the clusters was significantly decreased following MMF treatment compared to controls (Fig. 4D). This observation suggested that MMF treatment alters the molecular characteristics of Rac1 involved in the actin rearrangement and the dynamic behavior of microglial processes. Next, we categorized the subcellular location of the clusters into three primary localizations: the cell body, the origin of primary processes, and at the branching point of processes (Fig. 4E). Notably, as the majority of Rac1 clusters were found at the origins of the processes or the branching points (Fig. 4F), it is likely that these clustering structures are involved in process motility, particularly in process extension and retraction.

**Figure 4.**
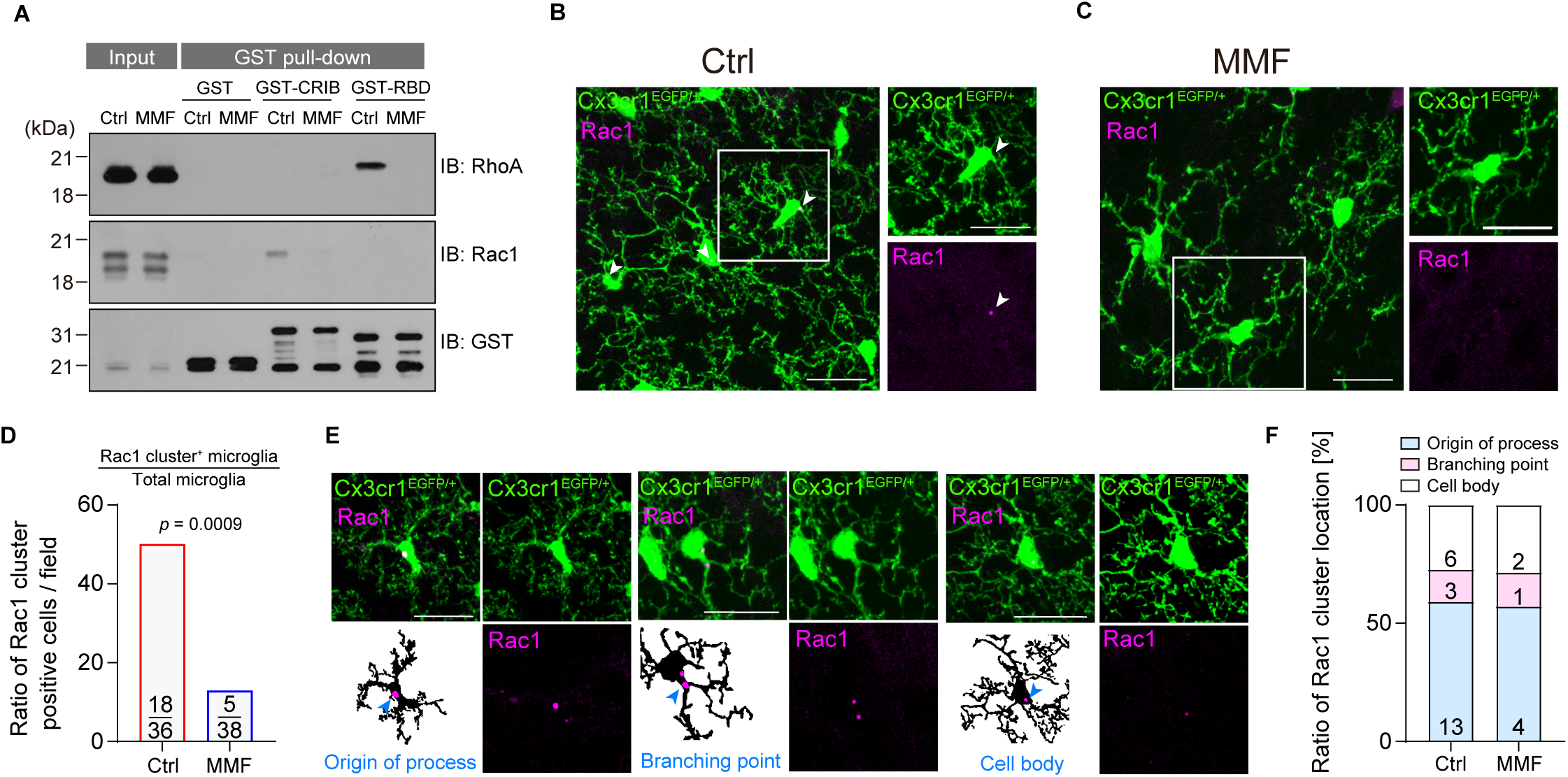
MMF treatment reduces the clustering of Rac1 in microglia. **(A)** GST pull-down assay using the mouse brain extraction (P14) with or without MMF administration. Precipitated samples were subjected to immunoblot analysis and to detect GTP-bound forms of RhoA and Rac1. **(B and C)** Immunostainings showing the expression of GFP and Rac1 in *Cx3cr1^GFP/+^* male mice with a vehicle control (B) or MMF (C). White arrows; Rac1 cluster. Scale bar = 20 µm. **(D)** Quantification of the ratio of Rac1 cluster positive microglia with vehicle-treated (Ctrl) or MMF-treated mice. Field = 1.03×10^5^ µm^2^. Ctrl ; number of microglial cells = 36, Rac1 cluster positive = 18. MMF ; number of microglial cells = 38, Rac1 cluster positive = 5. Fisher’s exact test. **(E)** Immunostainings showing the expression of GFP and Rac1 in *Cx3cr1^GFP/+^* male mice with a vehicle control. Rac1 cluster locate at the origin of process (1), branching point (2), or cell body (3). White arrows; Rac1 cluster. Scale bar = 20 µm. **(F)** The graphs show the percentage of the subcellular distribution of Rac1 clusters in (D). The number of clusters decreases, while the proportion remains unchanged.

### MMF treatment decreased the microglial complexity

Given that small GTPase proteins are involved in regulating morphological changes via actin rearrangement, we next analyzed the microglial morphology in the brain of *Cx3cr1^GFP/+^* mice with or without MMF treatment (Fig. 5A). First, we compared automated analysis of skeletonized microglia with manual quantification using ImageJ software (15,16). The results are consistent between the automated and manual analyses (Fig 5B and 5C). As it is challenging to quantify branching patterns using skeletonized automated quantification precisely, we analyzed the branching of their projections manually (Fig. 5D). Although there was no difference in the number of primary processes, the number of branching points arising from the primary processes was reduced after MMF administration (Fig 5E to 5H). On the other hand, the process lengths of MMF-treated microglia were consistent with those of wild-type microglia (Fig 5I to 5L). To further analyze the microglial complexity in the presence or absence of MMF, we performed a Sholl analysis (Fig 5M to 5N). We observed a significant decrease in the number of process intersections in MMF-treated microglia compared to controls, indicating a less complex cellular structure (Fig. 5O). Taken together, these data demonstrate that inhibition of IMPDH activity specifically impairs process branching, thereby attenuating the normal development of microglial morphological complexity in the early postnatal brain.

**Figure 5.**
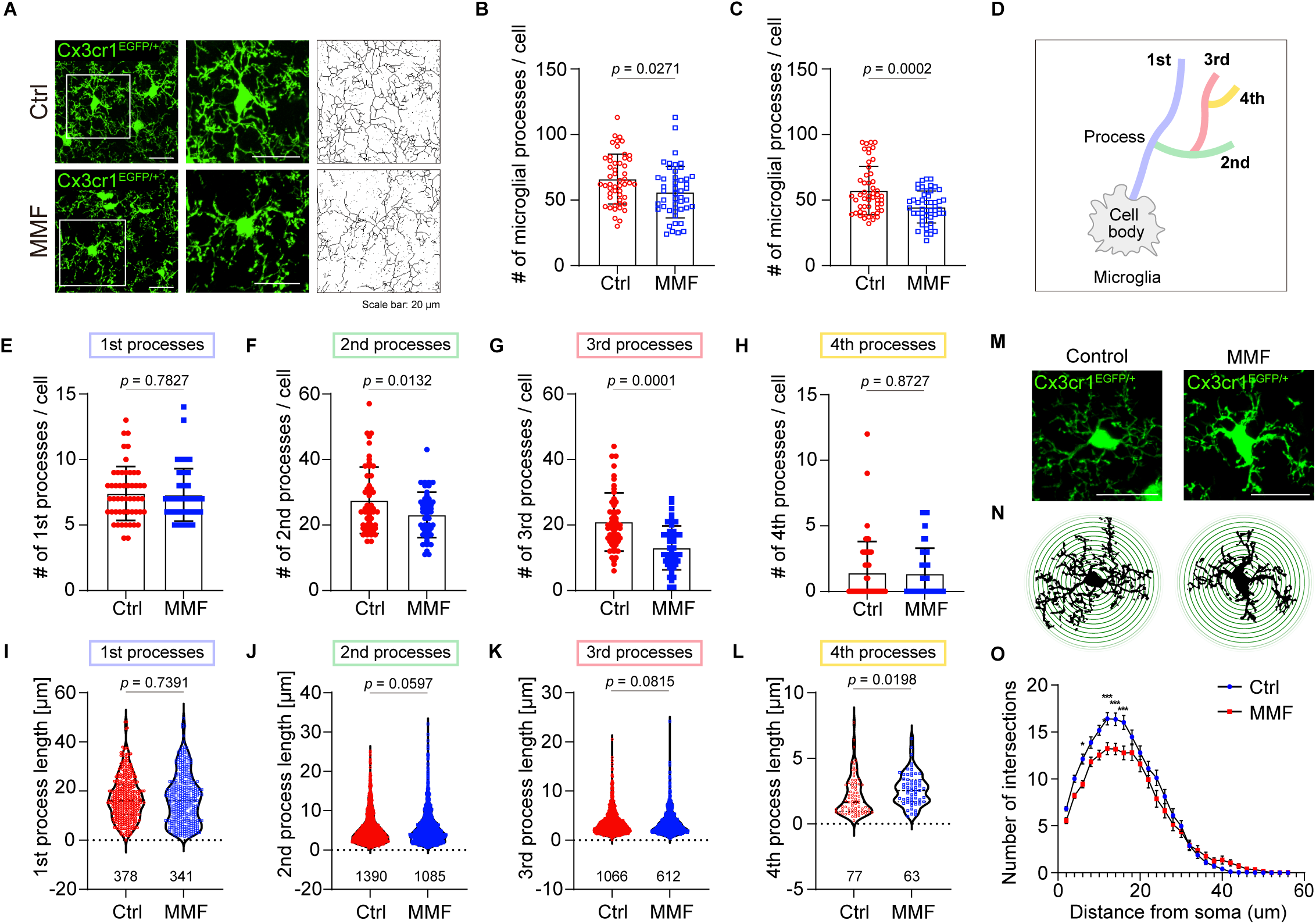
MMF treatment reduced the microglial complexity. **(A)** Immunostainings showing the expression of GFP in *Cx3cr1^GFP/+^* male mice. The right panel shows magnified insets and their corresponding skeletonized images. Scale bar = 20 µm. **(B and C)** Quantification of microglial processes using automated skeletonized assay (B) and manual assay (C). Ctrl; n = 3 mice, 51 cells, MMF; n = 3 mice, 47 cells. mean ± SEM, student’s *t*-test. **(D)** Schematic model of the microglial branching. **(E to H)** The manual quantification of the number of microglial processes in the cerebral cortex. (E) primary, (F) secondary, (G) tertiary, (H) quaternary processes. Ctrl; n = 3 mice, 51 cells, MMF; n = 3 mice, 47 cells. mean ± SEM, student’s *t*-test. **(I to L)** The manual quantification of the process length in cerebral cortex. (I) primary, (J) secondary, (K) tertiary, (L) quaternary processes. Ctrl; n = 3 mice, MMF; n = 3 mice. The number under the graph indicates the total number of cells counted. Mean ± SEM, student’s t-test. **(M)** Immunostainings showing the expression of GFP in *Cx3cr1^GFP/+^* male mice. Scale bar = 20 µm. **(N)** Schematic model of Sholl analysis in microglia with or without MMF. **(O)** Quantification of microglial intersections. Ctrl; n = 3 mice, 47 cells, MMF; n = 3 mice, 39 cells. mean ± SEM, student’s *t*-test.

## Discussion

In summary, our findings suggest that MMF administration plays a crucial role in regulating microglial branching, while not affecting the generation of primary processes or their elongation of them (Fig. 6). Since MMF administration decreases the level of guanosine nucleotides in the brain, the conversion of IMP to XMP may be a key process that regulates microglial properties during the early postnatal stages.

**Figure 6.**
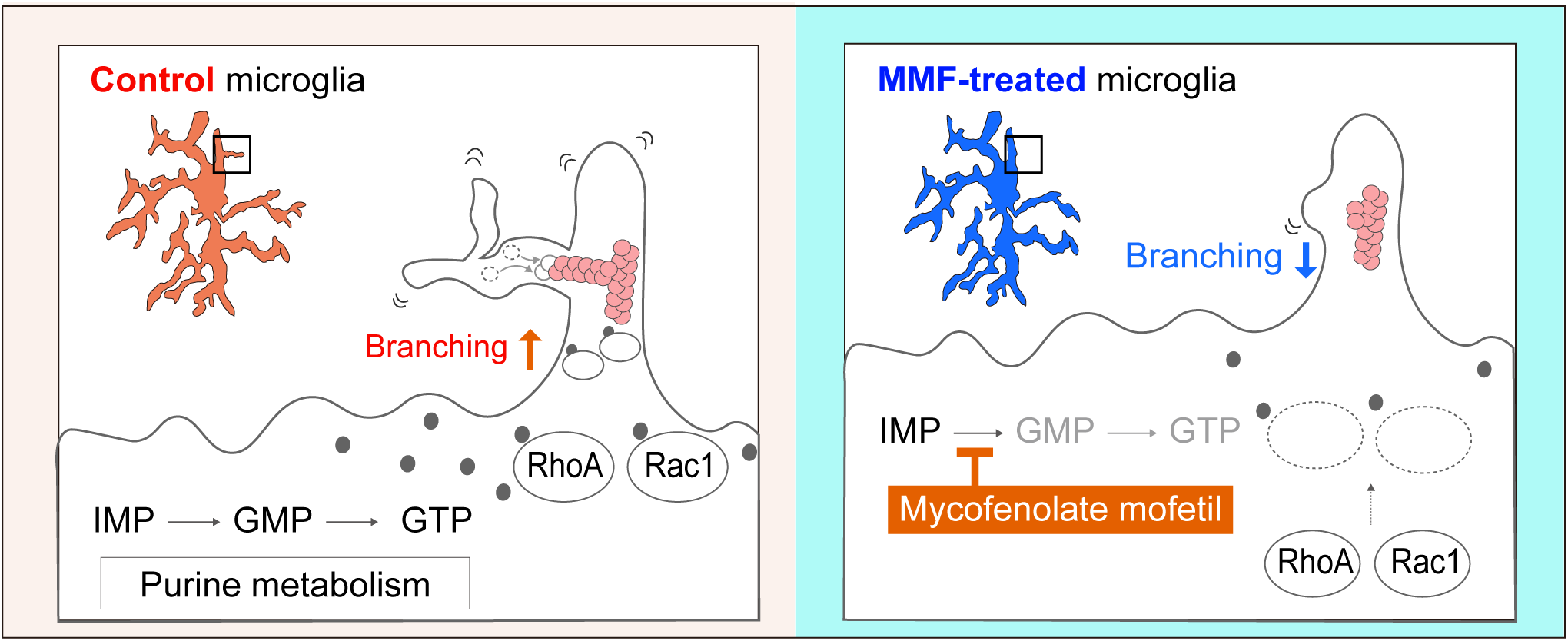
MMF treatment plays a crucial role in regulating microglial morphology. Model describing the effect of purine metabolism on the microglial morphology during the postnatal stage.

Previously it has been suggested that neural stem cells remain morphologically unchanged and are primarily influenced by MMF administration during mitosis (17). In contrast, we observed that microglial morphology is significantly altered when IMPDH activity is inhibited, although proliferation remains unaffected. These observations suggest that the activity of IMPDH may have distinct and cell-specific roles, indicating the complexity of purine metabolism in different cellular contexts. The expression of the purinosome complex, a master regulator for the de novo pathway, is initially high and subsequently stabilizes at a steady state as brain growth progresses. In contrast, the expression of HPRT gradually increases after birth (17). Notably, HPRT expression in microglia rises during the postnatal period, a time when gene expression patterns undergo changes in microglia (18). These observations suggest that the timing of the shift in purine metabolism from de novo to salvage pathways may be involved in the mechanisms by which IMPDH activity regulates cell growth and morphological changes.

We found that IMPDH inhibition reduces the level of GTP-bound small GTPase proteins (Rac1 and RhoA) and microglial complexity. This observation is consistent with previous work demonstrating that conditional knockout of *RhoA* gene in microglia results in a similarly simplified morphology, confirming that small GTPases activity plays an important role in this process (19). It is important to acknowledge, however, that GTPase activation is also dependent on the guanine nucleotide exchange factors (GEFs). While our data indicate that MMF reduces the available GTP pool, we cannot exclude the possibility that our findings reflect secondary effects on GEF activity or other signaling pathways. We also identified the formation of distinct Rac1 clusters within microglia, which are reminiscent of Cdc42 clusters in the previous study (14). While we show that the presence of these Rac1 clusters is significantly reduced by MMF administration, their precise function remains to be elucidated. Future studies using conditional knockout models and two-photon microscopy analysis will be useful to understand the exact contribution of these dynamic clusters to microglial morphology.

Our findings may also provide novel insight into neurodevelopmental disorders associated with purine metabolism, such as Lesch-Nyhan syndrome (LNS) (20,21). LNS is a rare genetic disorder caused by a deficiency in the salvage enzyme HPRT. This deficiency disrupts the recycling of purines, leading not only to systemic symptoms like hyperuricemia but also to severe neurological impairments such as self-injury. LNS patients are thought to compensate for the salvage pathway defect by upregulating the de novo synthesis pathway. This compensatory pathway renders IMPDH a key regulatory enzyme in the production of the guanosine nucleotides essential for brain development in LNS. On the other hand, based on our results, we propose a novel hypothesis that the neurological deficits observed in LNS arise from abnormal microglial development driven by insufficient IMPDH-dependent GMP synthesis during the critical early postnatal period.

Further investigations into IMPDH functions and their downstream effects on microglial transformation are expected to provide deeper insights into the mechanisms driving microglial morphogenesis and their functional implications in neurodevelopmental processes. Understanding these mechanisms may provide a potential cue for therapeutic strategies targeting microglial dysfunction in various developmental disorders.

## Materials and Methods

### Mice

The C57BL/6N mice were obtained from Japan SLC, Inc. (Shizuoka, Japan). The B6.129P2(Cg)-Cx3cr1^tm1Litt^ /J (005582) were obtained from the Jackson Laboratory. All male mice were maintained under a 12-hour light/12-hour dark cycle (lights on at 8:00 AM) in a temperature- and humidity-controlled environment. Mice were stimulated by MMF (60 mg/kg) intraperitoneally every 24 h from either P6 to P9 or P6 to P14.

### Immunohistochemistry

The *CX3CR1^GFP/+^* mice brains were perfused with PBS and fixed overnight in 4% paraformaldehyde (PFA) in PBS. After infiltration with 30% sucrose in PBS, the samples were embedded in Tissue-Tek OTC compound (Sakura Finetek, Tokyo, Japan) and sectioned at a thickness of 50 µm by a cryostat (CM1950, Leica Biosystems, Wetzlar, Germany). Free-floating sections were permeabilized and blocked with PBS containing 0.25% Triton X-100 and 5% bovine serum albumin (BSA). Primary antibodies were incubated in PBS containing 5% BSA for 1 day [chicken anti-GFP antibody (Abcam, Cat# ab13970, 1:1000) and mouse anti-Rac1 antibody (Bioscience, Cat#610650, 1:200)] at 4°C. Brain sections were washed with PBS and reacted with goat anti-chicken IgY (H+L) antibody, Alexa Fluor 488 (Thermo Fisher Cat# A11039, 1:1000) and donkey anti-mouse IgG (H+L) antibody, Alexa Flour 594 (Abcam, Cat# ab150108, 1:1000) and for 1 hour at room temperature. Samples were mounted in VECTASHIELD Mounting Medium (Vector Laboratories). Tissue specimens were observed using the confocal laser scanning microscope (LSM700, Carl Zeiss) with ×20 (Plan-Apochromat ×20/0.8 M27) and×63 (Plan-Apochromat ×63/1.4 Oil DIC M27) objectives. Diode excitation lasers (Diode 488) were operated and directed to a photomultiplier tube (LSM T-PMT, Carl Zeiss) through a series of bandpass filters. Z stack images (interval, 1 µm per image) were acquired using ZEN software (Carl Zeiss). The microglial skeletal assay and Sholl analysis were conducted using the ImageJ software. The manual measurements were performed by using the Freehand Tool in ImageJ.

### Imaging mass spectrometry

The mouse brains were frozen using dry ice quickly and stored -80 ℃. The brains were sectioned at a thickness of 10 µm using cryostat (CM1950, Leica Biosystems, Wetzlar, Germany) and were then put on Indium Tin Oxide coated glass slides (SI0100N, Matsunami, Osaka, Japan) under the -20 ℃ and stored at -80 ℃ with desiccant in 50 ml tubes. Matrix deposition was performed using iMLayer (Shimadzu, Kyoto, Japan). 9-aminoacridine (9-AA) (Merck KGaA, Darmstadt, Germany) was heated at 250 ℃ and deposited (0.9 µm) on the section mounted ITO glass slide. The 9-AA solution (4 mg/mL) in 70% ethanol was prepared and 500 µL of 9-AA solution was loaded into a 0.3 mm diameter airbrush (PS-270, GSI Creos, Tokyo, Japan) for additional coating via manual spray application, completing a two-step matrix coating. Matrix-assisted laser desorption/ionization mass spectrometry imaging (MALDI-MSI) was performed using IMScope QT (Shimadzu, Kyoto, Japan). MSI data were acquired in negative ion mode at an m/z range of 340–550. The detailed parameters are given in Supplementary Table 1. The MSI images were reconstructed using the data analysis software IMAGEREVEAL MS (Shimadzu, Kyoto, Japan).

### Purification of recombinant proteins

Escherichia coli BL21-Star (DE3) were transformed by pGEX6p-1, pGEX6p-1-RBD, pGEX6p-1-CRIB plasmids. Recombinant proteins were expressed in bacteria and purified using standard protocols. Bacterial cultures were induced with 1 mM isopropyl-β-D-thiogalactopyranoside (IPTG) at an optical density (OD_600_) of 0.5 and incubated at 37 °C for 6 hours. Cells were harvested and lysed by ultrasonication, and the lysates were clarified by centrifugation. The supernatants were applied to a glutathione Sepharose column (GE Healthcare) after filtration using 0.45 μm syringe filter, and bound proteins were eluted with 10 mM reduced glutathione. Purified recombinant proteins (GST, GST-RBD, and GST-CRIB) were dialyzed by PBS overnight at 4 °C.

### GST pull-down assay

For the pull-down from cell lysates, BV2 cells were plated at 2.0 x 10^6^ cells on 100 mm dish and incubated at 37°C with 5% CO_2_ for 1 day. BV2 cells were treated with or without 1 µM MMF in 2 hours and collected with 3 ml lysis buffer [20 mM Tris-HCl (pH 8.0), 150 mM NaCl, 1 mM EDTA, 0.5% NP-40, and 1 mM DTT]. Lysates were centrifuged at 14,000 rpm for 5 min, and supernatants were collected. The supernatants were mixed with 2 ug of each recombinant protein (GST, GST-RBD, and GST-CRIB) and glutathione-Sepharose beads (GE Healthcare) and incubated for 2 hours at 4 °C with rotation. The precipitants were washed 3 times with lysis buffer and subjected to immunoblot analysis. For the pull-down from the brain lysates, the mice with or without MMF administration were perfused with PBS and dissected using surgical scissors and forceps. The brains were transferred to microtubes and soaked in 1 ml of lysis buffer. The brains were homogenized using homogenizer pestle. Lysates were centrifuged at 14,000 rpm for 15 min, and supernatants were collected. The supernatants were mixed with 2 ug of each recombinant protein (GST, GST-RBD, and GST-CRIB) and glutathione-Sepharose beads (GE Healthcare) and incubated for 3 hours at 4 °C with rotation. The precipitants were washed 3 times with lysis buffer and subjected to immunoblot analysis.

### Immunoblot analysis

Samples were mixed with sample buffer (125 mM Tris-HCl pH 6.5, 4% SDS, 10% Glycerol, 0.01% bromophenol blue, 10% 2-mercaptoethanol) and boiled at 98°C for 3 minutes. After SDS-polyacrylamide gel electrophoresis, each sample was transferred to a polyvinylidene difluoride membrane (Pall Corp.). The membranes were blocked in 5% skim milk/TBST for 60 min at room temperature and were then incubated overnight at 4 °C with the primary antibodies [anti-Rac1 antibody (Bioscience, Cat#610650, 1:1000), anti-RhoA antibody (Santa Cruz, sc-418, 1:1000), and anti-GST antibody (Santa Cruz, B14, 1:1000)]. After washing with TBST, the proteins on membrane were detected with HRP-conjugated secondary antibodies (SeraCare, 1:20,000). The signals were detected by chemiluminescent substrate reagent (Chemi-Lumi One Super, Nacalai tesque)

### Time-lapse imaging of BV2 cells

BV2 cells were plated at 1.0 x 10^5^ cells on a 12-well plate and incubated for at 37°C with 5% CO_2_ for 1 day. Cells were pre-treated with MMF (1 µM) for 30 min prior to imaging analysis. Live cell imaging was conducted on fluorescence microscope (BZ-X800, KEYENCE) equipped with 10x Plan Apochromat 0.45 NA objective lenses (CFI Plan Apo λ10x, Nikon). Live-cell imaging was conducted by taking frames every 15 minutes for 135 minutes. Analysis of these images was performed in FIJI ImageJ, where the cell length (Feret diameter), cell area, and circularity were calculated for each cell through the "Set Measurements" function.

### Statistical analysis and image processing

The statistical analyses were calculated by using GraphPad Prism (GraphPad Software). All image processing was conducted by FIJI Image J 2.1.0/1.53c. The number of cells were quantified by Fiji/ImageJ software. No data were excluded from the statistical analysis. For comparisons between two groups, we used Student’s *t*-test. All representative microscopic images and western blotting shown in the figures reflect at least three biological replicates

## List of abbreviations

(EMPs): erythro-myeloid progenitors
(HGPRT): Hypoxanthine-guanine phosphoribosyl transferase
(MMF): mycophenolate mofetil
(IMPDH): inosine monophosphate dehydrogenase
(IMP): inosine monophosphate
(GMP): Guanosine monophosphate
(GDP): Guanosine diphosphate
(GTP): Guanosine triphosphate
(IMS): Imaging mass spectrometry
(GST): Glutathione S-transferase

## Acknowledgments

We thank Kseniya Sevostianova (University of Tsukuba) for editing the paper. We thank the lab members for helpful discussions and technical support.

## Authors’ contributions

R.T. and F.T. designed research; R.T., T.Y. performed research; R.T., K.U., T.T., T.Y., N.Y., H.K. analyzed data; R.T. and F.T. wrote the paper; F.T. Study supervision:

## Availability of data and materials

The datasets used in this manuscript are shown in the Source data file.

## Ethics approval and consent to participate

All experiments were approved by the animal experiment committee at the University of Tsukuba (the approval numbers: 23-370, 24-377, 25-365) and conducted according to the university guidelines for animal care.

## Consent for publication

Not applicable.

